# Targeting CyclinD1-CDK6 to Mitigate Senescence-Driven Inflammation and Age-Associated Functional Decline

**DOI:** 10.1101/2025.08.01.668243

**Authors:** Adarsh Rajesh, Aaron P. Havas, Rouven Arnold, Kathryn Lande, K. Garrett Evensen, Kelly Yichen Li, Sainath Mamde, Qian Yang, Armin Gandhi, Karl N. Miller, Marcos Garcia Teneche, Zoe Yao, Jessica Proulx, Andrew Davis, Laurence Haddadin, Michael Alcaraz, Carolina C. Macip, Brightany Li, Xue Lei, Charlene Miciano, Elizabeth Smoot, Allen Wang, Jeffrey H. Albrecht, April E. Williams, Bing Ren, Kevin Y. Yip, Peter D. Adams

## Abstract

Cellular senescence contributes to aging and age-related diseases by driving chronic inflammation through the Senescence Associated Secretory Phenotype (SASP) and interferon-stimulated genes (ISGs). Cyclin D1 (CCND1), a key cell cycle regulator, is paradoxically upregulated in these non-proliferating cells. We show that CCND1 and its kinase partner CDK6 drive SASP and ISG expression in senescent cells by promoting DNA damage accumulation. This leads to the formation of cytoplasmic chromatin fragments (CCFs) that activate pro-inflammatory CGAS-STING signaling. The tumor suppressor p53 (TP53) and its target p21 (CDKN2A) antagonize this CCND1-CDK6-dependent DNA damage accumulation pathway to suppress the SASP. In aged mouse livers, senescent hepatocytes show increased Ccnd1 expression. Hepatocyte-specific *Ccnd1* knockout or treatment with the Cdk4/6 inhibitor Palbociclib reduces DNA damage and ISGs in aged mouse liver. Notably, Palbociclib also suppresses frailty and improves physical performance of aged mice. These findings reveal a novel role for CCND1/CDK6 in regulating DNA damage and inflammation in senescence and aging, highlighting it as a promising therapeutic target.

## Introduction

Cellular senescence is a stable form of cell-cycle arrest triggered by stresses such as DNA damage, oncogene activation, and telomere shortening^1–7^. Senescent cells accumulate with age in many tissues and contribute to chronic inflammation, tissue dysfunction, and age-related pathologies through secretion of pro-inflammatory cytokines, chemokines, and interferon-stimulated genes (ISGs), collectively termed the senescence-associated secretory phenotype (SASP)^8–10^. Persistent DNA damage signaling in senescent cells promotes the formation of cytoplasmic chromatin fragments (CCFs) and activation of the cyclic GMP–AMP synthase (cGAS)–stimulator of interferon genes (STING) pathway, which sustains SASP and systemic inflammation^11–18^. Identifying the molecular drivers that maintain this chronic inflammatory state is essential for understanding and targeting age-related dysfunction.

Cyclin D1 (CCND1), classically defined as a regulator of G1 progression through activation of CDK4/6 and phosphorylation of the retinoblastoma protein (pRB) ^19–24^, is paradoxically elevated in senescence despite proliferative arrest^25–27^. The functional significance of CCND1 upregulation in non-proliferating, senescent cells remain unclear. Moreover, whether CCND1’s unconventional accumulation contributes causally to persistent DNA damage signaling, cytoplasmic chromatin stress, or inflammatory gene expression has not been explored.

Here, we investigate the role of CCND1 and its associated kinase CDK6 in sustaining DNA damage, cytosolic chromatin accumulation, and inflammatory signaling in senescence. Using complementary in vitro and in vivo models, we reveal an essential role for the CCND1–CDK6 complex in promoting persistent DNA damage, CCF formation, and cGAS–STING-driven inflammation. Mechanistically, we identify previously unrecognized interactions between CCND1 and chromatin-associated kinesin proteins, such as KIF4A, which has been implicated in chromatin architecture and DNA repair^28–31^. Finally, we show that genetic ablation of CCND1 in aged hepatocytes or pharmacological inhibition of CDK4/6 significantly attenuates chronic inflammatory signaling and ameliorates age-associated functional decline, suggesting broad therapeutic implications.

Collectively, our study identifies the CCND1–CDK6 axis as a novel mediator of senescence-associated inflammation and DNA damage stress, offering potential targets to mitigate age-related pathology and preserve tissue function in aging.

## Results

### CCND1 is expressed in senescent cells

Despite being a canonical driver of cell proliferation, prior studies have reported paradoxical elevations of cyclin D1 in senescent cells^25–27^. Consistent with these findings, we confirmed that CCND1 is upregulated at the RNA level in IMR90 fibroblasts following ionizing radiation (IR)-induced senescence. While most proliferation-associated genes were downregulated in senescence as expected, CCND1 was among a small group that showed consistent upregulation (Fig. 1A). A time course demonstrated progressive accumulation of cyclin D1 protein over 10 days after IR (Fig. 1B). Similar increases in CCND1 expression were observed by RNA-seq in previously published replication-induced and oncogene-induced senescence models^32^ (Fig. 1C). Western blotting confirmed that CCND1 and CCND2 are elevated in senescent IMR90 cells, accompanied by markers of senescence, downregulation of CCNA2, CCNB1, LMNB1 and phosphorylated pRB, and upregulation of CDKN1A and phosphorylated p65 NF-κB (Fig. 1D).

**Figure 1.**
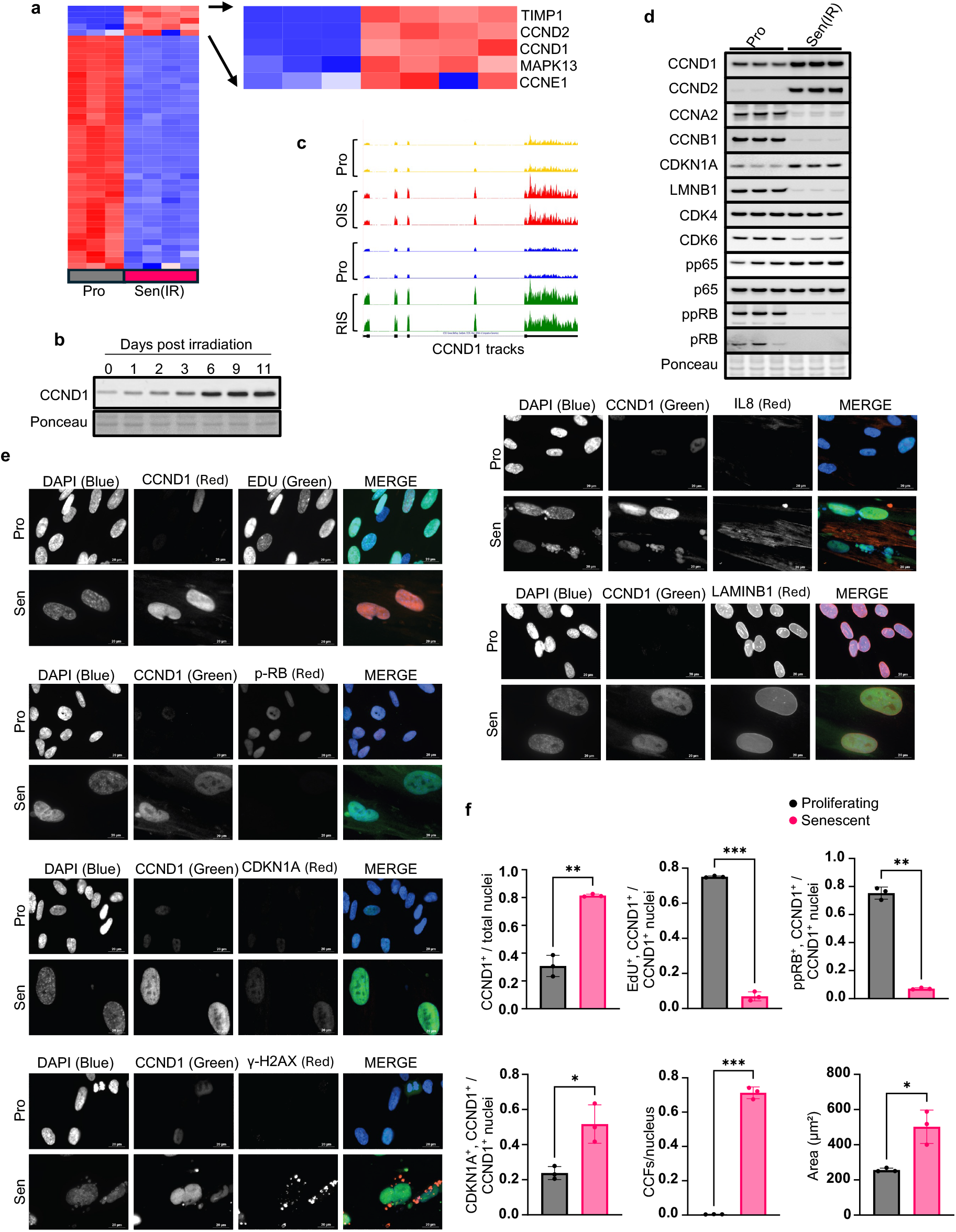
CCND1 is elevated in senescence and marks cells with canonical senescent features. a, RNA-seq of IMR90 fibroblasts 10 days after ionizing radiation (IR) shows CCND1 among a small subset of proliferation-associated genes upregulated in senescence. b, Western blot time course (0–11 days post-IR) shows progressive accumulation of cyclin D1 protein in senescent cells. Ponceau staining was used as a loading control c, Genome browser tracks show CCND1 upregulation in replication-induced and oncogene-induced senescence. d, Western blot of senescent IMR90s shows increased CCND1 and CCND2, decreased CDK6, and no change in CDK4. Phosphorylated pRB (ppRB) and total pRB were both reduced. Senescence markers CCNA2, CCNB1 and LMNB1 also decreased, while CDKN1A and phospho-p65 (pp65) were increased. Total p65 was unchanged. Ponceau staining was used as a loading control. **e**, Representative immunofluorescence images of proliferating (Pro) and senescent (Sen) IMR90s. In senescent cells, CCND1 localizes to nuclei that are EdU-negative, ppRB–negative, and express high CDKN1A and IL-8. CCND1+ cells also exhibit cytoplasmic chromatin fragments (CCFs; γH2AX/DAPI-positive puncta), enlarged nuclei, and reduced Lamin B1 at the nuclear periphery. f, Quantification of: percentage of total nuclei that are CCND1+; percentage of CCND1+ nuclei that are EdU+, ppRB+ or CDKN1A+; number of CCFs per nucleus; and nuclear area. Each dot represents an independent biological replicate (separate irradiation). For immunofluorescence quantifications, each dot is the average of ≥3 technical replicates from the same irradiation. Error bars denote mean ± s.d. Statistical analysis was performed using t-test. P < 0.05 was considered significant.

To confirm that CCND1-positive cells are senescent, we performed immunofluorescence staining on proliferating and IR-induced IMR90 fibroblasts (Fig. 1E, F). In senescent cells, cyclin D1 localized to EdU-negative and ppRB-negative nuclei and was co-expressed with CDKN1A, γH2AX and IL-8, a well-established SASP factor^25,33,34,6,7,9^. These CCND1-positive cells showed increased CCFs, defined as γH2AX/DAPI-positive chromatin in the cytoplasm, enlarged nuclei, a recognized feature of senescent hypertrophy^35–37^, and reduced Lamin B1 at the nuclear envelope^38,39^. Together, these features — cell cycle arrest, persistent DNA damage, inflammatory signaling, nuclear envelope disruption, CCF and cellular enlargement — indicate that elevated CCND1 marks cells exhibiting canonical molecular and structural hallmarks of senescence.

### CCND1 and CDK6 are required to sustain inflammatory and interferon gene expression in senescent cells

Expression of CCND1 in senescent cells is paradoxical not only because senescent cells have exited the cell cycle, but also because phosphorylation of pRB, the canonical CCND1 substrate, is undetectable in senescent cells (Fig. 1D). To investigate a potential function for this upregulation, we tested whether CCND1 and its kinase partners CDK4 and CDK6 (also expressed in senescent cells; Fig. 1B–D) are required to maintain senescence-associated transcriptional programs. We used two complementary approaches in IR-induced senescent IMR90 fibroblasts: siRNA-mediated knockdown of CCND1, CDK4 or CDK6, and pharmacological inhibition with Palbociclib (PD 0332991), a selective CDK4/6 inhibitor^40–43^ (Fig. 2A–B). Western blotting and qPCR confirmed efficient knockdown (Fig. 2C–D; Supplementary Fig. 1A). RNA-seq was performed 10 days after IR and PCA plots of transcriptomic data showed clear separation of CCND1/CDK6 knockdowns from non-targeting controls and of Palbociclib from DMSO-treated cells (Supplementary Fig. 1B–C). CCND1–CDK6 physical association in senescent cells was verified by CCND1 immunoprecipitation followed by Western blotting (Supplementary Fig. 1D).

**Figure 2.**
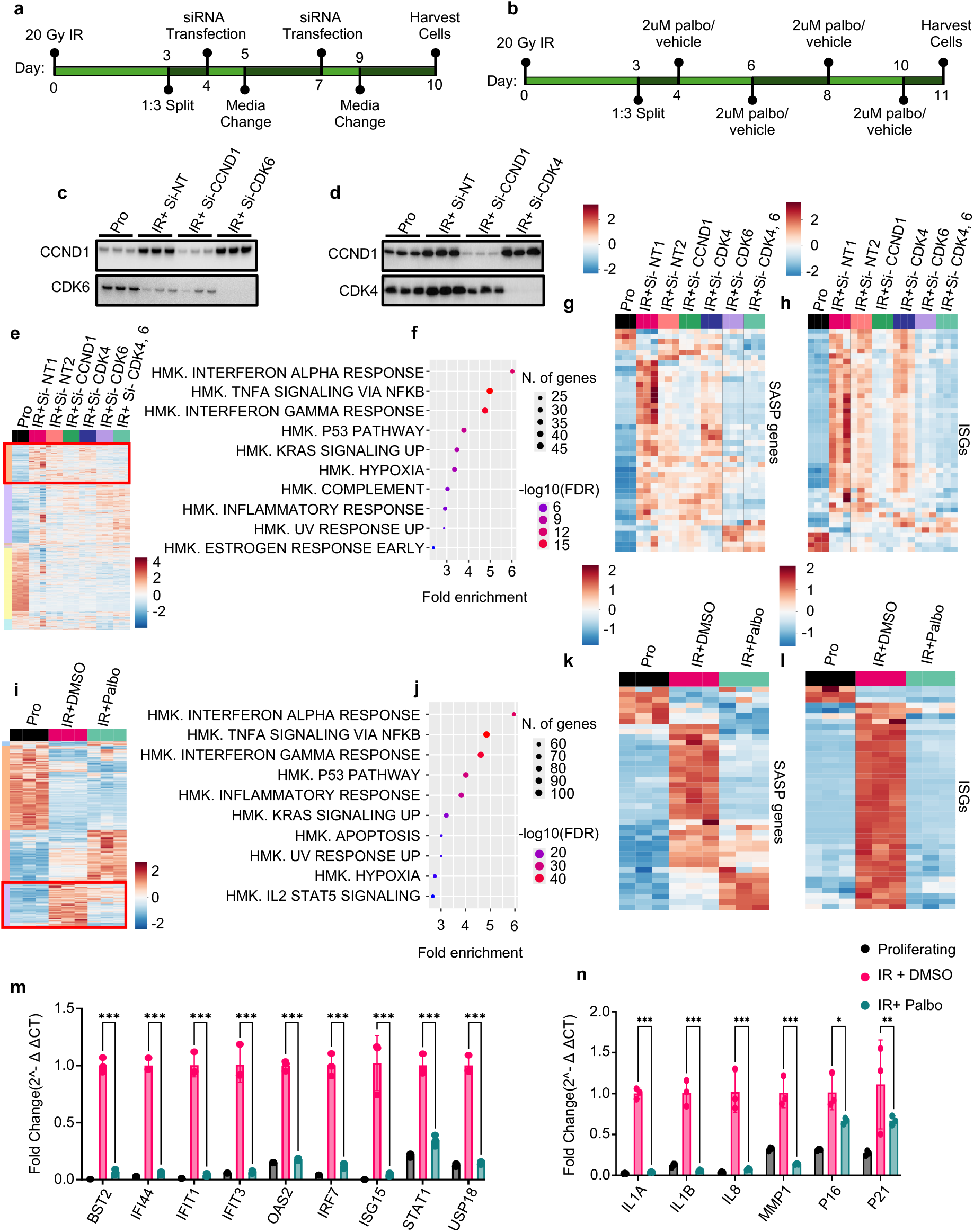
CCND1 and CDK6 are required to sustain inflammatory and interferon gene expression in senescence. a, Experimental design for siRNA knockdown experiment: IMR90 fibroblasts were induced senescent by ionizing radiation (IR) and transfected with siRNAs targeting CCND1, CDK4, CDK6, or both CDK4 and CDK6. Non-targeting controls (siNT1, siNT2) and proliferating (non-irradiated) cells were included. b, Experimental design for Palbociclib experiment: IMR90 fibroblasts were induced senescent by IR and treated with DMSO or Palbociclib. Proliferating (non-irradiated) controls were included. c–d, Western blots validating knockdown of CCND1, CDK4 and CDK6. e, Heatmap of all differentially expressed genes across siRNA conditions. A red box highlights a senescence-associated gene cluster selectively suppressed by CCND1 or CDK6 knockdown. f, Top enriched pathways in the red-box cluster include interferon-α response, TNFα signaling via NFκB, and interferon-γ response. g–h, Expression of SASP and ISG genes that are differentially expressed between proliferating cells and senescent siNT controls. i, Heatmap of all differentially expressed genes across Palbociclib conditions, with a red box indicating a cluster suppressed by Palbociclib treatment. j, Enrichment analysis of the Palbociclib-suppressed cluster shows reduced interferon-α response, TNFα signaling via NFκB, and interferon-γ response — the same top pathways as in f. k–l, Expression of SASP and ISG genes differentially expressed between proliferating and senescent DMSO-treated controls, showing suppression with Palbociclib. *m–n*, qPCR validation of SASP and ISG repression following Palbociclib treatment. Gene expression values are shown as fold change relative to senescent DMSO-treated controls, normalized to the geometric mean of GAPDH and RPL13. Each biological replicate represents an independent irradiation. Error bars denote mean ± s.d. Statistical analysis was performed using two-way ANOVA with Tukey’s post hoc test. *P* < 0.05 was considered significant.

Transcriptomic profiling following knockdown revealed broad changes in gene expression (Fig. 2E). Supplementary Fig. 2A–B shows a cluster of genes that increased upon CCND1 or CDK6 knockdown (and not CDK4), together with its hallmark enrichments. A distinct cluster of genes that were strongly upregulated in control senescent cells (NT1 and NT2) was selectively downregulated by CCND1 and/or CDK6 knockdown, but not by CDK4 knockdown (red box, Fig. 2E). Hallmark enrichment analysis of this CCND1/CDK6-dependent cluster indicated reduced interferon alpha and gamma responses, TNFα signaling via NFκB and p53 signaling after CCND1 and/or CDK6 knockdown (Fig. 2F). Heatmaps of curated SASP genes based on Coppé et al., 2010^44^ and ISGs based on Schoggins et al., 2011^45^ confirmed strong repression of inflammatory programs in CCND1- and/or CDK6-depleted cells, with minimal change after CDK4 knockdown (Fig. 2G–H).

Palbociclib (2 μM) was administered on days 4, 6, 8, and 10 post IR, following dosing regimens reported previously^46,47^, and recapitulated the transcriptional effects of CCND1 and CDK6 knockdown. Supplementary Fig. 2C–D presents the gene cluster increased by Palbociclib relative to DMSO and its enriched pathways. This FDA-approved drug suppressed a senescence-induced cluster analogous to the knockdown-sensitive cluster (red box, Fig. 2I), and hallmark analysis again identified reductions in interferon, inflammatory and p53 signaling (Fig. 2J). SASP and ISG heatmaps showed similar transcriptional repression after CDK4/6 inhibition (Fig. 2K–L). Consistent with these transcriptomic changes, Palbociclib reduced pSTAT1, total STAT1 and phospho p65 NFkB by Western blot, while total p65 was unchanged (Supplementary Fig. 1G). qPCR confirmed that representative SASP factors and ISGs were significantly downregulated by CCND1 and CDK6 knockdown (but not CDK4 knockdown) and by Palbociclib treatment (Supplementary Fig. 1E–F; Fig. 2M–N). Importantly, CCND1 or CDK6 knockdown or Palbociclib caused only a modest decrease in CDKN1A and CDKN2A. Moreover, cell cycle and proliferation-promoting genes were not upregulated by knock down (Supplementary Fig. 2C, 2F) confirming that CCND1 and/or CDK6 knock down did not cause cell cycle re-entry. Together, these findings suggest that CCND1 upregulation in senescent cells reflects an active role in sustaining pro-inflammatory and interferon-driven transcriptional programs, primarily through its kinase partner CDK6, and that these outputs can be suppressed via CCND1/CDK6 inhibition, without escape of cell cycle arrest.

### CCND1–CDK6 promotes DNA damage and CCF accumulation in senescent cells

To investigate how CCND1–CDK6 supports SASP and ISG expression in senescent cells, we tested whether it contributes to the accumulation of DNA damage and CCFs, precursors to SASP/ISG induction via cytosolic DNA signaling ^11–18^. Palbociclib significantly reduced DNA damage in IR-induced senescent IMR90s, as measured by Comet assay^48^ (Fig. 3A), and lowered 53BP1 and γH2AX protein levels on Western blot^49,50^ (Fig. 3B). Nuclear γH2AX intensity and CCF frequency also declined with Palbociclib (Fig. 3C). Consistent with these pharmacologic effects, siRNA knockdown of CCND1 or CDK6 similarly reduced nuclear γH2AX and CCFs (Supplementary Fig. 2E). In line with decreased CCF, Palbociclib-treated cells also showed decreased 2′3′-cGAMP by ELISA (Fig. 3D), indicating reduced cGAS–STING activation^13^.

**Figure 3.**
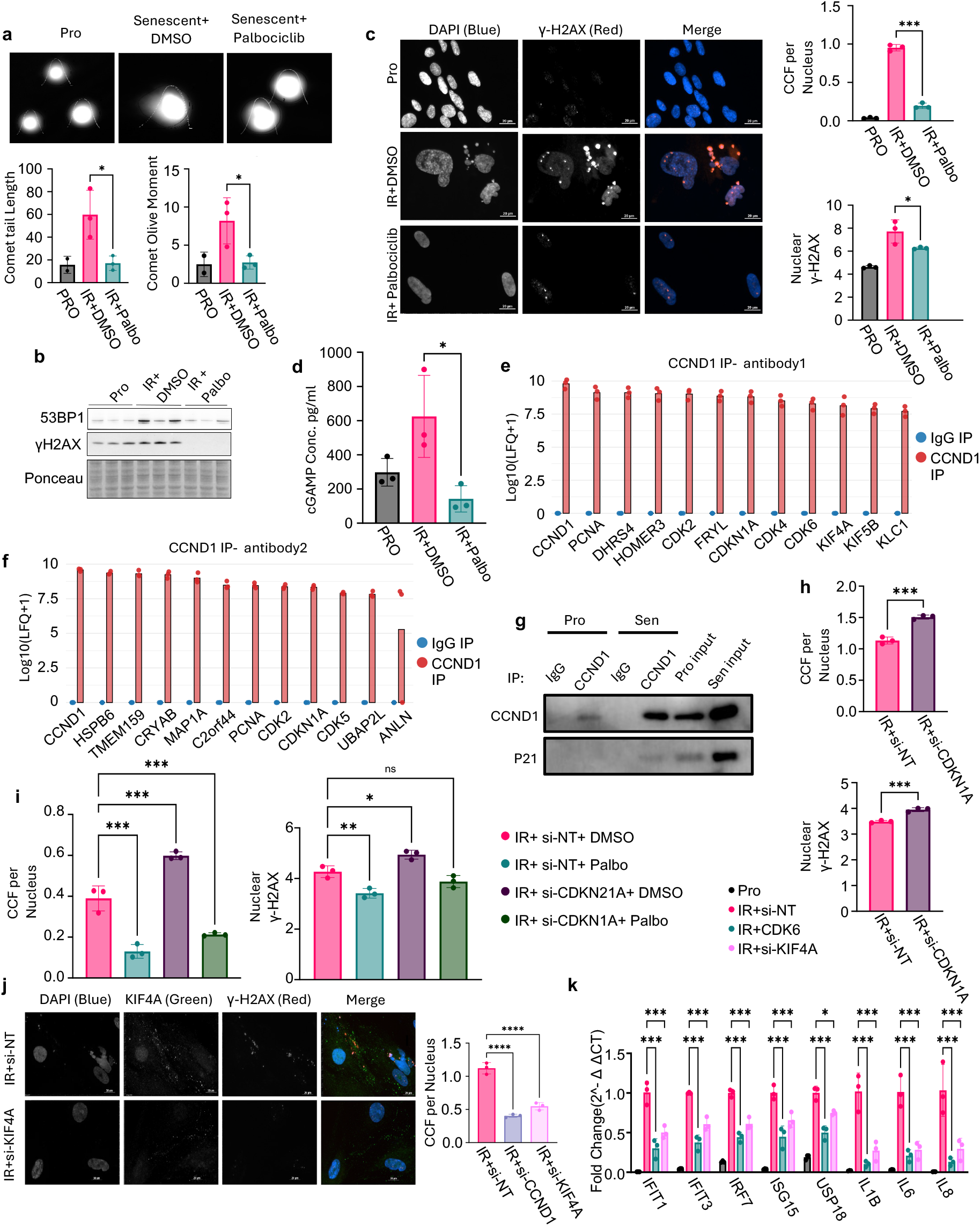
CCND1–CDK6 promotes DNA damage and cytoplasmic chromatin fragment (CCF) accumulation in senescent cells. a, Representative comet assay images and quantification of comet tail length and olive moment in IR-induced senescent IMR90 fibroblasts, showing reduced DNA damage with Palbociclib treatment. b, Western blots show decreased 53BP1 and γH2AX protein levels following Palbociclib; Ponceau staining was used as a loading control. c, Representative immunofluorescence images and quantification of nuclear γH2AX intensity and CCF frequency, both reduced in Palbociclib-treated senescent cells. d, ELISA for 2′3′-cGAMP shows reduced cGAS–STING activation after Palbociclib. e–f, Immunoprecipitation–mass spectrometry (IP–MS) using two CCND1 antibodies identifies top CCND1 interactors in senescent IMR90s, shown as ranked bar plots of log10(LFQ+1) intensity comparing D1 IP versus IgG control. g, Co-immunoprecipitation confirms interaction between CCND1 and CDKN1A in senescent cells. h, Knockdown of CDKN1A increases nuclear γH2AX and CCF formation. i, Palbociclib rescues the elevated DNA damage and CCFs caused by CDKN1A knockdown. j, Representative immunofluorescence images and quantification showing that KIF4A knockdown reduces CCF frequency. k, qPCR analysis showing that KIF4A knockdown suppresses SASP and ISG gene expression. Expression values are shown as fold change relative to non-targeting control siRNA in senescent cells, normalized to the geometric mean of GAPDH and RPL13. Each biological replicate represents an independent irradiation. For immunofluorescence quantifications, each dot reflects the average of ≥3 technical replicates per irradiation. Error bars denote mean ± s.d. Statistical analysis was performed using one-way ANOVA. *P* < 0.05 was considered significant.

CCND1 immunoprecipitation followed by mass spectrometry (IP-MS) using two distinct antibodies (rabbit and mouse) identified top-confidence interactors after IP of CCND1 in senescence compared to respective igG control IPs; their log10(LFQ+1) intensities are shown in Fig. 3E–F, with peptide counts and coverage in Supplementary Fig. 2F–G. We examined CDKN1A, PCNA and the kinesin proteins KIF4A, and KIF5B in more detail due to their known involvement in DNA damage response^28–31,51–63^, and prior data from our lab showing p53, upstream regulator of CDKN1A, to be a suppressor of DNA damage and CCF in senescent cells^63^. Co-immunoprecipitation confirmed the CCND1–CDKN1A interaction in senescent cells (Fig. 3G). CDKN1A knockdown increased nuclear γH2AX and CCFs (Fig. 3H) as we had reported previously^63^, and this phenotype were significantly reversed by Palbociclib (Fig. 3I). Together with the interaction between CCND1 and CDKN1A, this is consistent with a model whereby p53-CDKN1A restrains CCND1/CDK6–driven DNA damage and CCF formation. Among the kinesin proteins tested, only KIF4A knockdown phenocopied CCND1/CDK6 knockdowns, and reduced both CCFs (Fig. 3J) and SASP/ISG expression (Fig. 3K), implicating KIF4A as a candidate regulator and/or mediator of CCND1/CDK6-dependent chromatin stress and inflammation. Together, these data indicate that CCND1/CDK6 activity sustains DNA damage and cytosolic chromatin stress in senescent cells, thereby maintaining inflammatory gene expression. This function is likely restrained by p53-CDKN1A and mediated, at least in part, through chromatin-associated factors such as KIF4A.

### CCND1 is elevated in aged hepatocytes and promotes DNA damage and inflammation

Aged liver has been shown to accumulate senescent cells^64–67^. To investigate the functional role of potentially elevated Ccnd1 in aged livers, we first assessed its expression in hepatocytes from young and old mice. qPCR revealed a significant age-associated increase in Ccnd1 transcript levels (Fig. 4A). Immunofluorescence staining showed increased CCND1 protein in hepatocytes from old livers, specifically in Ki67-negative cells, suggesting accumulation in non-proliferating hepatocytes (Fig. 4B). An old mouse with a spontaneous liver tumor was included as a proliferative control, showing strong Ki67 positivity. Supporting this observation, immunohistochemistry confirmed increased CCND1 protein in livers of old mice, with signal restricted to phospho–Histone H3–negative cells, indicating non-dividing hepatocytes (Supplementary Fig. 3A). Partial hepatectomy liver samples served as positive controls for mitotic cells.

**Figure 4.**
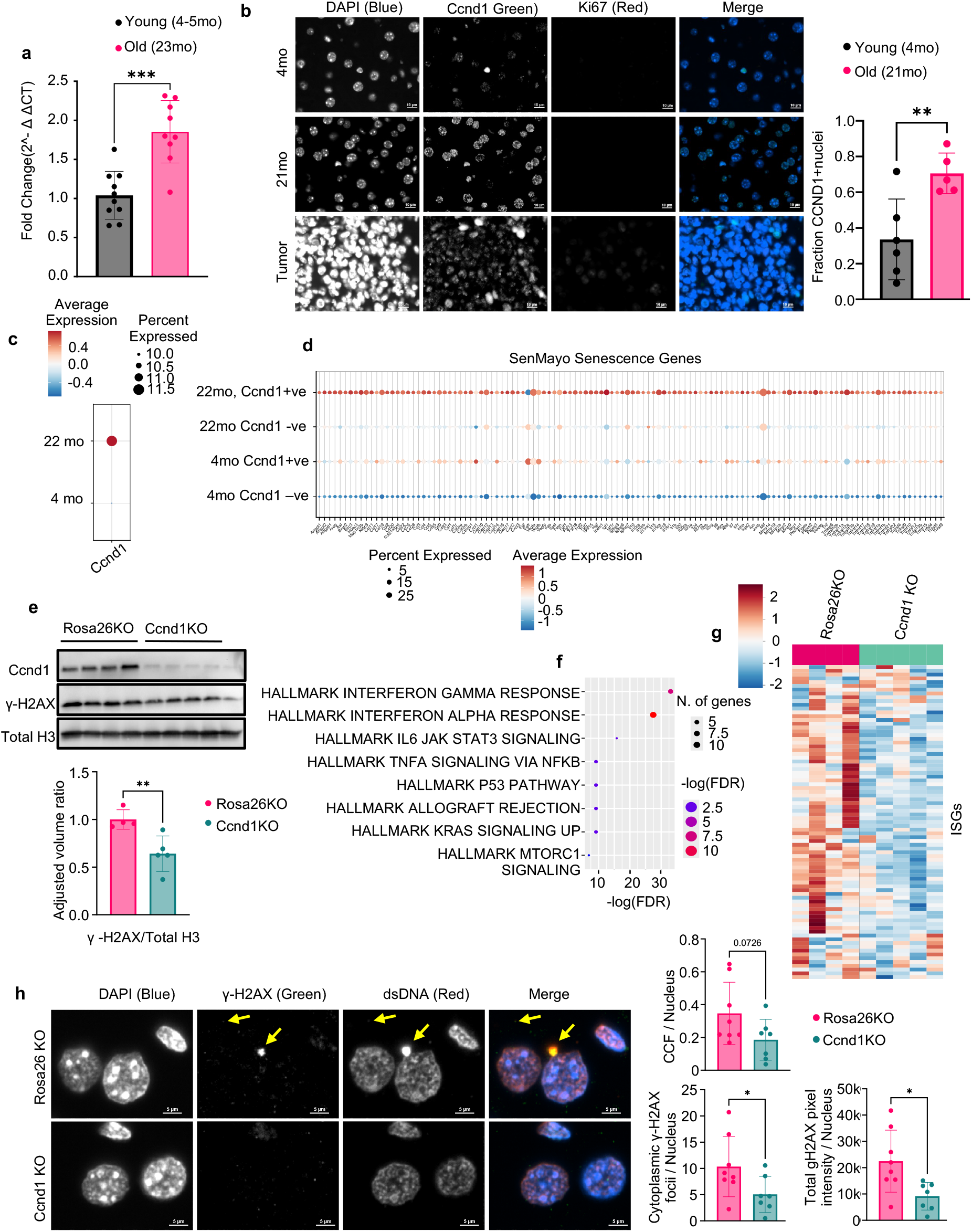
CCND1 is elevated in aged hepatocytes and drives DNA damage and inflammation in vivo. a, qPCR analysis of isolated hepatocytes from young (4–5-month-old) and old (23-month-old) mice showing increased *Ccnd1* expression with age. Expression values are shown as fold change relative to young and normalized to the geometric mean of GAPDH and HPRT. b, Immunofluorescence staining and quantification of liver sections from young (4-month-old) and old (21-month-old) mice showing increased CCND1 expression in hepatocyte nuclei of aged livers. CCND1 signal was observed predominantly in Ki67-negative cells, indicating non-dividing hepatocytes. c, CosMx spatial transcriptomics of livers from young (4-month-old) and old (22-month-old) mice showing elevated *Ccnd1* transcript levels in hepatocytes of aged livers. d, CosMx spatial transcriptomics showing that *Ccnd1*-positive hepatocytes in 22-month-old livers are enriched for SenMayo senescence-associated genes compared to *Ccnd1*-negative hepatocytes. e, Western blot analysis of liver lysates from 17-month-old mice 3 weeks after hepatocyte-specific *Ccnd1* knockout using AAV8–TBG–saCas9, showing marked reduction in CCND1 protein levels and γH2AX, confirming knockout efficiency and decreased DNA damage; Ponceau staining was used as a loading control. f, Bulk RNA-seq of liver tissue from 22-month-old mice showing broad suppression of inflammatory gene sets following *Ccnd1* knockout. g, Heatmap showing downregulation of interferon-stimulated genes (ISGs) that are typically elevated with age in livers of 22-month-old mice. h, Immunofluorescence analysis of liver sections from 22-month-old mice showing reduced cytoplasmic γH2AX puncta and a trend toward fewer cytoplasmic chromatin fragments (CCFs; overlapping γH2AX- and dsDNA-positive puncta in the cytoplasm, example shown with arrow) in *Ccnd1*-knockout livers. Each biological replicate represents a separate animal. Error bars denote mean ± s.d. Statistical analysis was performed using Welch’s t-test. For qPCR comparisons, Mann–Whitney U test was used. *P < 0*.*05 was considered significant*.

To assess whether Ccnd1-expressing hepatocytes exhibit features of senescence in vivo, we performed spatial transcriptomics (CosMx) on livers from young (4-month) and old (22-month) mice^68^. Ccnd1 transcript levels increased with age in hepatocytes (Fig. 4C), and in aged livers, Ccnd1-positive hepatocytes were enriched for the SenMayo senescence gene signature (Fig. 4D) ^69^, along with elevated expression of Cdkn1a and interferon-stimulated genes (ISGs) (Supplementary Fig. 4A– B), both of which are associated with senescence. To further confirm this association, we analyzed single-cell RNA sequencing data generated by the SenNet consortium from an independent cohort of young (3–4 months) and old (21–23 months) male and female mice. Consistent with the CosMx findings, Ccnd1+ hepatocytes displayed strong enrichment for both the SenMayo and ISG senescence signatures compared to Ccnd1− cells in both age groups (Supplementary Fig. 4C–F). Together, these data suggest that Ccnd1 expression marks hepatocytes with a senescent transcriptional phenotype in vivo.

To test whether Ccnd1 in aged hepatocytes is also functionally required for the increased inflammatory signaling reported with age^63–65^, we performed hepatocyte-specific Ccnd1 knockout using AAV8-mediated delivery of a saCas9 and sgRNA under the TBG promoter^70–74^ in 17-month-old mice. After three weeks, Ccnd1 was knocked out at both the DNA (Supplementary Fig. 3B) and protein levels (Fig. 4E). γH2AX levels were also decreased significantly with Ccnd1 knockout (Fig. 4E). Transcriptomic profiling revealed broad suppression of inflammatory pathways (Fig. 4F), including downregulation of ISGs that we previously observed to be elevated in the aged liver^74^ (Fig. 4G). These findings suggest that CCND1 promotes age-associated inflammatory gene expression in hepatocytes.

A 3-month-long Ccnd1 knockout was performed in young (3-month) and old (22-month) mice to assess whether these effects persist long term. Interferon-stimulated gene (ISG) transcripts were significantly reduced in aged Ccnd1 knockout livers compared to age-matched Rosa26KO controls^75^ (Supplementary Fig. 3C). At the protein level, CCND1 remained depleted at endpoint, and levels of γH2AX, STAT1, and phosphorylated STAT1 were also decreased as assessed by Western blot (Supplementary Fig. 3D).

Immunofluorescence also showed a trend toward reduced CCFs, defined as cytoplasmic puncta with overlapping dsDNA and γH2AX signals, along with a significant reduction in cytoplasmic γH2AX foci (Fig. 4H). Antibody specificity for γH2AX in CCF detection was validated by increased nuclear staining in irradiated young liver compared with non-irradiated controls (Supplementary Fig. 3E). Together, these findings indicate that age-associated Ccnd1 expression in hepatocytes marks senescent-like cells and is functionally required to sustain DNA damage and inflammatory gene expression in the aged liver.

### CDK4/6 inhibition reduces inflammation and protects against age-associated functional decline

To assess whether CDK4/6 inhibition can suppress chronic inflammation and improve physiological function in aged mice, we treated animals with the CDK4/6 inhibitor Palbociclib (64 mg/kg) via oral gavage for 3 weeks in two independent cohorts. In the first cohort, daily dosing was initiated in 8-month-old and 25-month-old mice. In the second cohort, dosing was administered three times per week (Monday, Wednesday, Friday) to 6-month-old and 23-month-old mice. In both experiments, Western blot analysis of spleen tissue confirmed target engagement, as evidenced by reduced levels of phosphorylated RB (ppRB) in Palbociclib-treated mice (Fig. 5A). qPCR analysis of liver tissue revealed a consistent trend toward reduced expression of interferon-stimulated genes (ISGs) in Palbociclib-treated mice compared to vehicle-treated controls in both dosing regimens (Fig. 5B–C).

**Figure 5.**
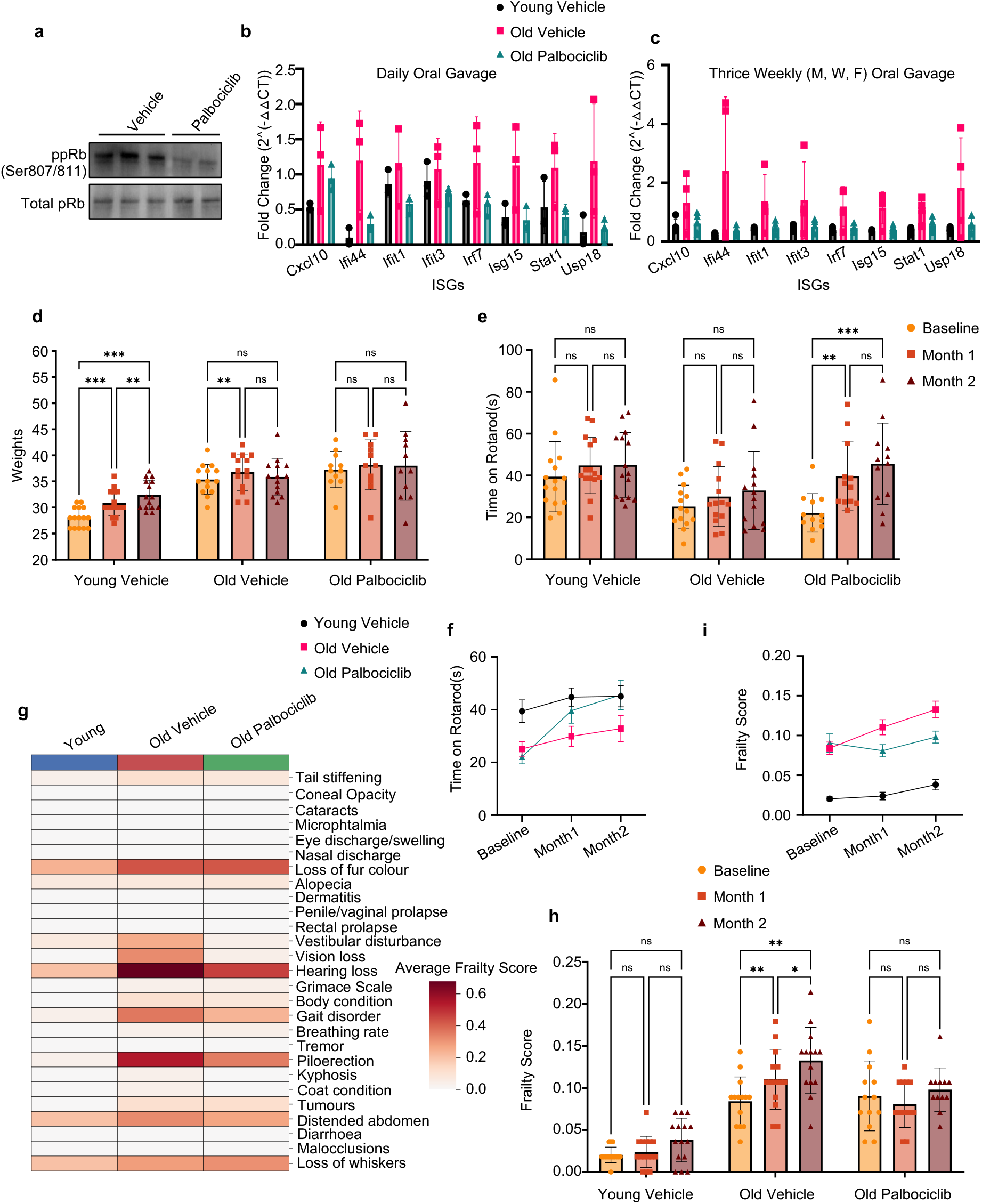
CDK4/6 inhibition reduces inflammatory gene expression and preserves function in aged mice. a, Western blot of spleen lysates from aged mice treated with Palbociclib, showing reduced phosphorylated pRB (ppRB), confirming target engagement; Ponceau staining was used as a loading control. b–c, qPCR analysis of liver tissue from two short-term Palbociclib dosing cohorts. Mice received 64 mg/kg Palbociclib in autoclaved H_²_O via oral gavage (200 μL) either daily (aged 25 months, young controls 8 months) or three times per week (aged 23 months, young controls 6 months) for 3 weeks. Both regimens showed a trend toward reduced interferon-stimulated gene (ISG) expression in Palbociclib-treated aged mice relative to age-matched controls. Expression values are shown as fold change relative to aged vehicle-treated mice and normalized to the geometric mean of GAPDH and HPRT. d, Body weight over the course of the long-term treatment study, showing no significant difference between Palbociclib- and vehicle-treated aged mice. e, Rotarod test performance at baseline and after two months of Palbociclib treatment in 18-month-old mice, demonstrating improved motor coordination and endurance in treated animals, comparable to 4-month-old young controls. f, Longitudinal rotarod trajectories over the two-month treatment course showing progressive improvement in Palbociclib-treated aged mice compared vehicle-treated aged mice. g, Subcomponent analysis of frailty scores showing the strongest Palbociclib-associated improvements in gait, hearing, vestibular function, and vision. h, Frailty index at the end of the study: Palbociclib-treated aged mice maintained stable frailty scores, whereas vehicle-treated aged mice showed increased frailty over time. i, Longitudinal frailty index trajectories showing divergence between vehicle and Palbociclib-treated groups across the treatment period. Each biological replicate represents a separate animal. Error bars denote mean ± SEM. Statistical analysis for longitudinal and paired measurements was performed using two-way repeated measures ANOVA. *P* < 0.05 was considered significant.

To evaluate long-term functional outcomes, 18-month-old mice were treated with Palbociclib (64 mg/kg, 3×/week) for two months and assessed longitudinally for motor coordination and frailty^76,77^. No differences in body weight were observed between groups during treatment (Fig. 5D). Rotarod performance progressively improved in Palbociclib-treated aged mice, and by the end of treatment their latency to fall was indistinguishable from that of 4-month-old controls, while age-matched vehicle-treated mice showed no improvement (Fig. 5E). These performance differences were also captured as average performance trajectories over time (Fig. 5F).

The protective effect of Palbociclib was most pronounced in specific frailty domains, including gait disorders, hearing loss, vestibular disturbance, and vision impairment (Fig. 5G). Overall frailty scores increased significantly over the two-month period in vehicle-treated aged mice but remained stable in the Palbociclib-treated group (Fig. 5H), indicating that CDK4/6 inhibition prevents worsening of frailty. Longitudinal analysis confirmed that average frailty scores diverged progressively between the two aged groups (Fig. 5I). Together, these results demonstrate that CDK4/6 inhibition can reduce inflammatory gene expression and protect against age-associated motor and physiological decline in mice.

## Discussion

Our findings reveal a previously unappreciated role for the canonical cell-cycle regulator CCND1 in cellular senescence and age-associated inflammation. Classically recognized as a driver of proliferation through its association with CDK4/6, the paradoxical upregulation of CCND1 during senescence suggests a fundamental shift in function. Rather than supporting proliferation, CCND1– CDK6 activity emerges as a critical determinant of sustained DNA damage signaling, cytoplasmic chromatin accumulation, and persistent inflammatory responses associated with non-proliferative senescent phenotypes.

CCND1 accumulation was consistently observed across multiple senescence models, including DNA damage-induced, replicative, and oncogene-induced senescence, despite global suppression of proliferative programs. While canonical regulators such as Cyclin A2, Cyclin B1, Lamin B1, and phosphorylated pRB were downregulated, CCND1 was among a small set of factors selectively upregulated. Its co-expression with senescence markers including γH2AX, IL-8, and CDKN1A, alongside features such as nuclear envelope disruption and CCFs, identifies CCND1 as a *bona fide* feature of senescence.

Mechanistically, our data position CCND1–CDK6 activity as essential for maintaining the pro-inflammatory transcriptional landscape of senescence, particularly SASP and ISG expression. Notably, CDK6, but not CDK4 was required to sustain these programs, highlighting a functional divergence between CDK4/6 complexes in senescent cells. Both genetic depletion of CCND1 or CDK6 and pharmacological inhibition via Palbociclib led to reductions in nuclear γH2AX, CCF formation, and cGAMP production, ultimately suppressing downstream inflammatory gene expression. These results suggest that CCND1–CDK6–driven DNA damage fuels cGAS–STING activation, thereby reinforcing chronic inflammation. This role is distinct from their well-established pro-proliferative functions and defines a previously unrecognized activity of CCND1–CDK6 in amplifying DNA damage–induced innate immune signaling during senescence.

Notably, our mass spectrometry analysis revealed that CCND1 interacts with a set of chromatin-associated proteins in senescent cells, including KIF4A and KIF5B. KIF4A has been implicated in chromatin compaction and DNA repair foci formation, suggesting a potential role in modulating chromatin stress through an interaction with CCND1–CDK6. While previous studies described KIF4A as facilitating DNA repair^28^, our data indicates a context-specific function in senescent cells, where KIF4A appears to support sustained DNA damage and inflammatory signaling. This divergence may reflect altered repair dynamics in senescent versus proliferating cells. KIF5B, another CCND1 interactor, has recently been implicated in the formation of nuclear envelope tubules that capture DNA double-strand breaks, thereby promoting their repair^31^, suggesting CCND1–CDK6 may also influence chromatin architecture and immune activation through kinesin-mediated transport mechanisms. Future experiments should dissect the functional contribution of kinesins and their physical interactions with CCND1–CDK6 to better understand their role in sustaining senescence-associated DNA damage and inflammatory signaling.

Importantly, our findings identify CDKN1A as a critical negative regulator of this pathway. Although CDKN1A expression is a hallmark of senescence-associated growth arrest, its depletion paradoxically exacerbates DNA damage and CCF formation, effects that are reversed by Palbociclib treatment. We have previously shown that p53, an upstream regulator of CDKN1A, also suppresses DNA damage and CCF formation in senescent cells^63^. Together, these results suggest that the p53–CDKN1A axis functions to counterbalance CCND1–CDK6–mediated genotoxic stress, coordinating pro-inflammatory signaling with cell cycle arrest. How CDKN1A mechanistically constrains CCND1–CDK6 activity, whether through direct inhibition of kinase activity, modulation of chromatin structure, or disruption of protein–protein interactions, remains an important question for future investigation.

In vivo, we observed marked CCND1 accumulation in aged, non-proliferative hepatocytes, aligning with our in vitro data and further supporting its role in driving senescence-associated phenotypes. Spatial transcriptomics revealed that CCND1-positive hepatocytes in aged mice were enriched for SenMayo genes, elevated ISGs, and high Cdkn1a expression, indicating a strong senescent signature. Functional ablation of CCND1 in hepatocytes led to reduced γH2AX, suppressed interferon signaling, and decreased cytoplasmic chromatin accumulation, confirming that CCND1 is not merely a marker but an active promoter of inflammatory stress in aged liver. These observations validate our in vitro findings and demonstrate that CCND1’s senescence-promoting functions extend beyond cultured fibroblasts to mammalian tissues.

From a translational perspective, systemic CDK4/6 inhibition in aged mice reduced inflammation and conferred notable protection against age-related functional decline. Palbociclib treatment reversed declining motor coordination and prevented progression of frailty, effects associated with reduced interferon signaling in vivo. These outcomes reinforce the therapeutic potential of targeting CCND1– CDK6-driven senescence mechanisms to mitigate inflammaging and extend healthspan. Given the existing clinical applications of CDK4/6 inhibitors for cancer^42,43,78–80^, their repositioning for age-related inflammatory diseases appears promising and warrants further exploration.

In summary, our study uncovers a novel and unexpected pro-senescence function of CCND1–CDK6 complexes that is intricately linked to promotion of DNA damage, innate immune signaling, and chronic inflammation during aging. These findings fundamentally revise our understanding of CCND1 biology and senescence regulation, highlighting previously unrecognized opportunities for therapeutic intervention in aging and age-related diseases. Future studies should explore the precise molecular mechanisms underlying CCND1-mediated chromatin dynamics, its interactions with kinesin-associated partners, and the broader implications of CCND1–CDK6 axis modulation across multiple tissues and aging contexts.

While CDK4/6 inhibitors like Palbociclib are widely used to induce senescence in cancer cells, this is often accompanied by a robust SASP, including pro-inflammatory cytokines and interferon responses^81–83^. In contrast, our study focuses on non-transformed cells, where we find that CDK6 activity sustains the SASP, and its inhibition suppresses chronic inflammatory signaling. Supporting this distinction, it has been shown that CDK4/6 inhibition in normal proliferating cells can trigger a senescence-like arrest with a blunted SASP^84^, consistent with our observations. These findings underscore the context-dependent effects of CDK4/6 inhibition and suggest that targeting the CCND1–CDK6 axis in aging, non-cancerous tissues may suppress inflammation, highlighting its potential as a therapeutic strategy for mitigating age-related inflammatory decline.

## Methods

### Animals

All animal procedures were approved by the Institutional Animal Care and Use Committee (IACUC) of Sanford Burnham Prebys Medical Discovery Institute. Animal experiments were performed at the Sanford Burnham Prebys Animal Facility in compliance with IACUC guidelines and all relevant ethical regulations for animal research. Young and old C57BL/6J mice were obtained from the NIA aging colony housed at Charles River Laboratories. “Young” mice were defined as 4–8 months of age, and “old” mice ranged from 18 to 25 months at the time of tissue collection.

### Palbociclib treatment in mice

Palbociclib isethionate (PD-0332991 isethionate, CAS 827022-33-3; eNovation Chemicals) was dissolved in autoclaved distilled water and administered by oral gavage in a total volume of 200 µl per mouse. Mice were dosed at 64 mg/kg, calculated based on an average body weight of 35 g. All animals received the same dose regardless of individual weight. For short-term studies, 23-25 month-old male mice received Palbociclib either daily or 3 times per week for 3 weeks. For long-term functional studies, 18-month-old male mice were treated 3 times per week for 2 months and evaluated for rotarod performance and frailty index alongside age-matched vehicle-treated controls and 4-month-old young controls. Frailty assessments were performed blinded.

### SaCas9 sgRNA design and validation

Guide RNAs were designed using CRISPick (Broad Institute) with the mouse reference genome GRCm38 (Ensembl v102). The sgRNA targeting Ccnd1 (forward: 5′-CACCGTGACACCAATCTCCTCAACGA-3′; reverse: 5′-AAACCTCGTTGAGGAGATTGGTGTCAC-3′) was annealed and cloned into the vector pX602-AAV-TBG::NLS-SaCas9-NLS-HA-OLLAS-bGHpA;U6::BsaI-sgRNA, as described previously^74^. Plasmids were transformed into NEB Stable competent *E. coli*, and successful insertion was confirmed by Sanger sequencing (Genewiz) using the universal U6 primer (5′-GACTATCATATGCTTACCGT-3′). To assess indel formation in mouse liver, genomic DNA was extracted using the Monarch Genomic DNA Purification Kit (NEB), and PCR was performed using primers flanking the target site (~500 bp amplicon). Products were purified with the QIAquick PCR Purification Kit (Qiagen) and submitted for Sanger sequencing using the forward amplification primer. Indel frequency was quantified using ICE analysis (Synthego ICE v3.0).

### AAV generation and usage

AAV8 particles were generated and purified as previously described^74^. Briefly, large-scale production was performed by the Salk Viral Vector Core or the SBP Functional Genomics Core. HEK-293T cells were co-transfected with the AAV vector plasmid, RepCap8, and Helper plasmid (Cell Biolabs) using polyethylenimine (Polysciences). After 96 hours, virus was harvested from both cell pellets and supernatants. Supernatants were precipitated using polyethylene glycol, and cell pellets were lysed by sonication. Crude lysates were treated with Benzonase (Millipore Sigma), and AAV particles were purified by iodixanol gradient ultracentrifugation. The fraction containing purified rAAV8 was collected, buffer-exchanged with sterile PBS, aliquoted, and stored at –80 °C. Viral titers were determined by SYBR Green qPCR using vector-specific primers.

### AAV-mediated hepatocyte-specific Ccnd1 knockout

Recombinant AAV8–TBG–SaCas9-U6-sgCcnd1 or AAV8–TBG–SaCas9-U6-sgRosa26 (control) was administered via retro-orbital injection at a dose of 3 × 10^11^ viral genomes per mouse in a 100 μL volume. The sgRNA targeting *Ccnd1* was validated as described previously^74^. Mice were harvested either 3 weeks or 3 months post-injection for short- and long-term experiments, respectively. Liver tissue was either flash-frozen for RNA and protein analysis or fixed for histology and immunofluorescence staining.

### Hepatocyte Isolation

As previously described^74^, hepatocyte isolation was performed following two-step perfusion. Mice were euthanized by CO_²_ asphyxiation and surgically exposed. A 24G × ¾″ catheter (Surflo, SR-0×2419CA) was inserted into the inferior vena cava below the liver. The liver was perfused with 50 mL of Hank’s Balanced Salt Solution (HBSS, Gibco, Ref 14175-095) pre-warmed to 42 °C at a flow rate of 6 mL/min, followed by 40 mL of DMEM (Gibco, Ref 10313-021) containing Collagenase Type IV (Gibco, Ref 17104-019), also pre-warmed to 42 °C. After digestion, livers were excised and disassociated in ice-cold DMEM with 2% Bovine Serum Albumin (BioWorld, CAS: 9048-46-8), then stored on ice. Cell suspensions were centrifuged at 50 × g for 2 min at 4 °C and washed five times in cold DMEM to remove debris. The resulting cell pellet was resuspended in a 40% Percoll solution (Cytiva, Ref 17089102), composed of 10% 10× HBSS (Gibco, Ref 14065-056) and 60% DMEM, then centrifuged at 100 × g for 7 min at 4 °C to enrich viable hepatocytes. The resulting pellet was washed once more with DMEM at 50 × g and resuspended. Viability and cell counts were determined using a hemocytometer and trypan blue staining.

### Liver Tissue Processing and Protein Extraction

As previously described^74^, liver tissue samples were flash-frozen in liquid nitrogen and stored at – 80 °C until processing. For protein extraction, tissues were homogenized either in RIPA buffer (Thermo Fisher) supplemented with protease and phosphatase inhibitors (Roche), or directly in 1× Laemmli buffer using a Precellys homogenizer. Homogenates were centrifuged at 10,000 × g for 15 minutes at 4 °C, and supernatants were collected. Protein concentrations were determined using the Bradford assay (Pierce, Cat# 1863028) and measured on a Spectra Max 190 plate reader. Equal amounts of protein were separated by SDS-PAGE using the Bio-Rad Mini-PROTEAN Tetra System and transferred to PVDF membranes (Millipore, Cat# ISEQ00010) at 100 V for 70 minutes. Membranes were stained briefly with Ponceau S (Sigma-Aldrich, Cat# P7170-1L) to confirm transfer, then blocked in 5% milk (BD Difco, Cat# 232100) in TBST for 1 hour at room temperature. Primary antibodies were diluted in 5% BSA (BioWorld, Cat# 9048-46-8) in TBST and incubated overnight at 4 °C on a rocker. The following primary antibodies were used: Cyclin D1 (Thermo Fisher Scientific, Cat# MA5-16356), Phospho-Histone H2A.X (Ser139) (Cell Signaling Technology, Cat# 9718S), Phospho-Stat1 (Ser727) (Cell Signaling Technology, Cat# 9177S), and Stat1 (Cell Signaling Technology, Cat# 9172S). Secondary HRP-conjugated antibodies included Goat anti-Mouse IgG-HRP (Thermo Fisher Scientific, Cat# 31446, RRID: AB_228318) and Goat anti-Rabbit IgG-HRP (Millipore, Cat# AP307P, RRID: AB_92641). Imaging was performed on a Bio-Rad ChemiDoc Touch Imaging System and analyzed using Image Lab software.

### Immunofluorescence – Liver Tissue

As previously described^74^, liver tissue was fixed in 10% Neutral Buffered Formalin (Epredia, Ref 9400-1) at room temperature for 48 hours, then paraffin-embedded and sectioned at 5 μm thickness. Tissue sections were deparaffinized with two sequential washes in 100% xylene, followed by 95%, 90%, and 70% ethanol. Antigen retrieval was performed using 10 mM Tris, 1 mM EDTA, and 0.05% Tween-20 in a steamer (Nesco, ST-25F) for 40 minutes. Slides were permeabilized for 15 minutes in 0.05% Triton X-100 in PBS and then blocked for 2 hours at room temperature in 4% BSA and 1% FBS. Primary antibodies were applied overnight at 4 °C and included Cyclin D1 (Thermo Fisher Scientific, Cat# MA5-16356), γH2AX (Histone H2A.X Ser139ph; Active Motif, Cat# 39118), and Ki-67 (SolA15 clone; Thermo Fisher Scientific, Cat# 14-5698-82). After washing, sections were incubated with fluorophore-conjugated secondary antibodies for 1 hour at room temperature. Autofluorescence was quenched using 0.1% Sudan Black (Sigma, Cat# 199664) in 70% ethanol for 1 hour. DAPI was applied at a concentration of 5 μg/mL for 15 minutes. Coverslips were mounted using Fluoromount-G (Thermo Fisher Scientific, Ref 00-4958-02), and images were captured using a Nikon T2 fluorescence microscope.

### Cell Culture and Senescence Induction

IMR90 primary human lung fibroblasts (ATCC CCL-186) were cultured as described previously^63^ in Dulbecco’s Modified Eagle Medium (DMEM; Gibco 10313-121) supplemented with 10% fetal bovine serum (FBS; Corning 35-010-CV), 1% penicillin-streptomycin (Gibco 15140-122), and 2 mM glutamine (Gibco 25030-081). Cells were maintained at 37 °C in a humidified incubator with 5% CO_²_ and 3.5% O_²_. All cell lines tested negative for mycoplasma contamination and were used at passages 12–25. Senescence was induced by exposing 20–30% confluent cultures to 20 Gy X-ray irradiation using a Cs-137 source. Cells were allowed to reach confluence over 3 days and subsequently passaged for downstream experiments.

### siRNA Transfection

Senescent IMR90 fibroblasts were transfected with 50 nM siRNA (Dharmacon siGENOME SMARTpool) using 0.8% DharmaFECT 1 reagent according to the manufacturer’s instructions. For most knockdown experiments, transfections were performed on day 4 and day 7 after irradiation. Cells were returned to complete media 18–20 hours after each transfection and harvested on day 10 post-IR. For siP21, a single transfection was performed on day 4, and cells were collected on day 7. A pool of four siRNAs was used per gene. All experiments were performed in biological triplicate.

### Palbociclib Treatment (In Vitro)

Palbociclib hydrochloride (Selleckchem, Cat# S1116) was dissolved in DMSO to a stock concentration of 10 mM and stored at –20 °C. Senescent IMR90 fibroblasts were treated with 2 μM Palbociclib starting on day 4 post-irradiation. Treatments were administered on days 4, 6, 8, and 10 post-IR. Cells were harvested on day 11. Control cells were treated with an equivalent volume of DMSO vehicle on the same days. Treatments were refreshed by media replacement at each time point.

### Comet assay

DNA damage was assessed using the neutral comet assay following the manufacturer’s instructions (Enzo Life Sciences, Cat# ADI-900-166). Briefly, cells were embedded in low-melting-point agarose on slides and incubated at 4°C in the dark for 30 minutes. Slides were then immersed in pre-chilled lysis buffer (2.5 M NaCl, 100 mM EDTA pH 10, 10 mM Tris base, 1% sodium lauryl sarcosinate, 1% Triton X-100) for 45 minutes at 4°C. Electrophoresis was carried out in TAE buffer (40 mM Tris base, 20 mM acetic acid, 1 mM EDTA) for 15 minutes at 30 V at room temperature. After electrophoresis, slides were washed and stained according to kit protocol. Comet tails were quantified using an open-source ImageJ plugin^85^.

### cGAMP ELISA

Cyclic GMP–AMP (cGAMP) levels were quantified using a competitive ELISA kit (Cayman Chemical, Cat# 501700) according to the manufacturer’s instructions. Liver tissues were homogenized in M-PER Mammalian Protein Extraction Reagent (Thermo Fisher) and clarified by centrifugation. Protein concentration was measured using the Bradford assay (Pierce), and equal amounts of lysate were used for the ELISA. Absorbance was read using a microplate reader, and cGAMP concentrations were interpolated from a standard curve of known cGAMP standards provided by the kit.

### Quantitative RT–PCR

RNA was isolated using TRIzol (Invitrogen, Ref 15596018) according to the manufacturer’s recommendations. For liver tissues, RNA was purified using the Zymo Direct-zol RNA Miniprep Kit (Zymo Research). For cultured cells, RNA was extracted using chloroform separation followed by isopropanol precipitation. RNA concentrations were measured using a Nanodrop One spectrophotometer (Thermo Scientific). cDNA synthesis was performed using RevertAid Reverse Transcriptase (Thermo Fisher, Ref EP0441), Ribolock RNase Inhibitor (Thermo Fisher, Ref EO0381), and 5× RT Reaction Buffer (Thermo Fisher). Gene expression was quantified using PowerUp SYBR Green Master Mix (Applied Biosystems, Ref A25741) on a QuantStudio 6 Flex Real-Time PCR System (Applied Biosystems, Ref 4485691).

### Western Blotting (Cell Culture)

For in-vitro experiments, IMR90 cell lysates were initially prepared in RIPA buffer (Thermo Fisher) supplemented with Halt protease and phosphatase inhibitor cocktail (Thermo Fisher Scientific), then denatured in NuPAGE LDS sample buffer (Thermo Fisher, Ref# NP0007) with reducing agent. Proteins were resolved on NuPAGE 4–12% Bis-Tris gels (Thermo Fisher Scientific) using MES SDS running buffer at 140 V and transferred to PVDF membranes (Immobilon PSQ, Millipore, Ref# ISEQ00010) using the Bio-Rad Trans-Blot Turbo semidry transfer system for 55 minutes. Membranes were blocked in 5% milk in TBST, incubated overnight at 4 °C with primary antibodies diluted in 5% BSA, followed by HRP-conjugated secondary antibodies. Blots were imaged using a Bio-Rad ChemiDoc Touch imaging system, and band intensities were quantified using Image Lab software. The following primary antibodies were used: Cyclin D1 (Thermo Fisher Scientific, Cat# MA5-16356), Cyclin D2 (Cell Signaling Technology, Cat# 3741S), Cyclin A2 (Abcam, Cat# ab38), Cyclin B1 (Millipore, Cat# 4220), CDKN1A/p21 (Santa Cruz Biotechnology, Cat# sc-817), Lamin B1 (ProteinTech, Cat# 12987-1-AP), CDK4 (Cell Signaling Technology, Clone D93GE), CDK6 (Cell Signaling Technology, Cat# 13331), pp65 (Ser536) (Cell Signaling Technology, Cat# 3033S), p65 (Santa Cruz Biotechnology, Cat# sc-8008), STAT1 (Cell Signaling Technology, Cat# 9172S), phospho-STAT1 (Ser727) (Cell Signaling Technology, Cat# 9177S), pRb (Cell Signaling Technology, Cat# 9313S), ppRb (Ser807/811) (Cell Signaling Technology, Cat# 8516S), phospho-Histone H2A.X (Ser139) (Cell Signaling Technology, Cat# 9718S), and 53BP1 (Millipore, Cat# MAB3802). Secondary HRP-conjugated antibodies included Goat anti-Mouse IgG-HRP (Thermo Fisher Scientific, Cat# 31446, RRID: AB_228318) and Goat anti-Rabbit IgG-HRP (Millipore, Cat# AP307P, RRID: AB_92641).

### Immunofluorescence – Cell Culture

Cells were plated on PhenoPlate™ 96-well microplates (PerkinElmer), stained as described previously^86^. Following fixation, permeabilization, and blocking, cells were incubated overnight at 4 °C with primary antibodies. The following antibodies were used: Cyclin D1 (Thermo Fisher Scientific, Cat# MA5-16356 and Santa Cruz Biotechnology, Cat# sc-20044), ppRB (Ser807/811; Cell Signaling Technology, Cat# 8516S), CDKN1A/p21 (Santa Cruz Biotechnology, Cat# sc-817), γH2AX (Histone H2A.X Ser139ph; Active Motif, Cat# 39118 or Millipore, Cat# 05-636), IL-8 (Abcam, Cat# ab18672), Lamin B1 (ProteinTech, Cat# 12987-1-AP), and KIF4A (Thermo Fisher Scientific, Cat# PA5-30492). After washing, appropriate fluorophore-conjugated secondary antibodies were applied for 1 hour at room temperature. Nuclei were counterstained with DAPI (5 μg/mL). Plates were imaged using a Nikon T2 fluorescence microscope with automated image capture. Image analysis was performed in NIS Elements AR v5.21.03 using dark background subtraction, thresholding, size exclusion, and automated partitioning to quantify signal.

### Immunoprecipitation and Mass Spectrometry (IP-MS)

Immunoprecipitation was performed as previously described^87^ with the following modifications. Cells were lysed in EBC100 buffer (50 mM Tris-HCl pH 8.0, 100 mM NaCl, 0.5% NP-40) supplemented with protease and phosphatase inhibitors (Roche), rotated at 4 °C for 30 minutes, and cleared by centrifugation at 12,000 × g for 10 minutes. Protein concentration was determined using the Bradford assay (Pierce). Dynabeads™ Protein G (Thermo Scientific, 10004D) were conjugated to one of two primary antibodies against CCND1: Rabbit monoclonal Cyclin D1 antibody (Cell Signaling Technology #2922S) and Mouse monoclonal Cyclin D1 antibody (DCS-6) (Santa Cruz Biotechnology, sc-20044). Beads were pre-bound to antibody for 45 minutes at room temperature, then incubated with lysate overnight at 4 °C. Control IPs were performed using corresponding rabbit or mouse IgG. The following day, beads were washed with cold NETN buffer (20 mM Tris pH 8.0, 1 mM EDTA, 100 mM NaCl, 0.5% NP-40) five times for western blot (IP-WB) or three times with NETN followed by two washes with TBS for mass spectrometry (IP-MS). A portion of the eluate (10%) was retained for western blot validation and the remainder was snap-frozen. IP-MS was conducted by the Proteomics Core at SBP Medical Discovery Institute. Bead-bound proteins were digested using trypsin/Lys-C after denaturation, reduction, and alkylation, and peptides were desalted using C18 cartridges. LC-MS/MS analysis was performed using an EASY-nanoLC coupled to a Q-Exactive Plus Orbitrap with a 120-minute gradient and standard DDA settings as previously described.

### RNA Sequencing and Analysis

PolyA-enriched RNA libraries were prepared using either the Watchmaker Genomics mRNA Library Prep Kit (Watchmaker Genomics) with Elevate Long UDI Adapters (Element Biosciences), or the NEBNext Poly(A) mRNA Magnetic Isolation Module followed by the NEBNext Ultra II Directional RNA Library Prep Kit (NEB). Sequencing was performed as 2×75 bp paired-end reads on either the Element Biosciences AVITI sequencer (Cloudbreak 2×75 kit) or the Illumina NextSeq500. Library preparation method and sequencing platform are noted in figure legends or supplementary data as applicable. Adapter and quality trimming was performed using Trim Galore (v0.6.7) or Trimmomatic (v0.38.1). Reads were aligned to the human genome (hg38) or mouse genome (mm10) using HISAT2 (v2.2.1). Gene-level counts were quantified using featureCounts (v2.0.3). Quality control was performed using FastQC (v0.74) and summarized with MultiQC. Differential expression analysis and normalization were conducted using DESeq2 (v2.11.40.8), and principal component analysis (PCA) was used to assess sample clustering and relationships.

### 10x Single-Cell Sequencing

Raw sequencing reads were processed using cellranger arc (v2.0.2). Ambient RNAs were removed using CellBender (v0.3.0) [Cite: PMID 37550580]. Cells marked as doublets by DoubletFinder (2.0.4) [Cite: PMID 30954475] were removed. Additional data processing was performed using Seurat (v5.2.1) [Cite: PMID 37231261] and Signac (v1.12.0) [Cite: PMID 34725479]. We also discarded cells with >10% mitochondrial reads, <200 genes detected, <1000 ATAC reads, or ATAC enrichment at TSS < 1. The samples were then merged, normalized using SCTransform and integrated using Harmony. After cell clustering by Seurat, initial cell type annotation for each cluster was performed based on canonical marker genes. For each initial cell type, we performed subclustering and checked the expression of marker genes for each subclusters. Subclusters expressing multiple marker genes were removed. Hepatocytes were considered Ccnd1+ if their normalized expression of Ccnd1 was greater than 0. For comparing expression of SenMayo genes and ISGs between Ccnd1-positive and Ccnd1-negative hepatocytes, only genes expressed in more than 10% of hepatocytes were kept.

### CosMx Spatial Transcriptomics

Flat files for each CosMx slide were processed using Seurat (v4.9.9.9050). The individual tissue samples from each slide were split and to be treated as individual samples. The FOVs for each tissue (nine/tissue) were combined to form one large FOV/tissue. Cells in the top and bottom 15th percentile for number of features and with counts less than 20 were removed. The samples were then re-merged, normalized using SCTransform (v0.3.5) with a clip range of −10 to 10, and integrated using Harmony (v1.2.3). Each cluster, as determined by Seurat, was further subclustered (resolution 0.1 to 0.3). Cell type annotation was first checked using CellKB (https://www.cellkb.com)^92^ after filtering for only “liver” studies. The top five markers, sorted by log2 fold change were used as input genes for CellKB. Cell type annotation was further confirmed using known markers for liver cell types (Immune Cells: Ptprc; B Cells: Cd79a; Cholangiocytes: Spp1, Krt19, Epcam, Sox9; Hepatocytes: Apoa1, Glul, Serpina1a, Apoe; HSC: Dcn, Bmp5, Col1a1, Col1a2; Kupffer Cells: Cd5l, Cd74, C1qa, C1qb; LSEC: Lyve1; NK Cells: Stat4, Nkg7; T Cells: Cd2, Cd3d, Cd3g). Hepatocytes were considered Ccnd1+ if their normalized expression of Ccnd1 was greater than 0.

## Data Availability

All RNA-seq and spatial transcriptomic datasets generated in this study have been deposited in the Gene Expression Omnibus (GEO) under the following accession numbers: GSE304160: CosMx spatial transcriptomics of young and old mouse liver, GSE304265: Bulk RNA-seq of liver tissue from hepatocyte-specific *Ccnd1* knockout and Rosa26 control mice, GSE304266: Bulk RNA-seq of IMR90 fibroblasts following irradiation and siRNA knockdown of *CCND1* or *CDK6*, GSE304272: Bulk RNA-seq of irradiated IMR90 fibroblasts treated with Palbociclib.

## Acknowledgments

This work was supported by the National Institutes of Health under grants P01 AG031862, U54 AG079758 (SenNet San Diego Tissue Mapping Center), P01 AG073084, and K99 AG073450, as well as by the American Society of Hematology (ASH) and the CIRM Training Grant EDUC-4-12813. We thank Adriana Charbono and Buddy Charbono for technical support with mouse experiments, and Guillermina Garcia and Monica Sevilla from the histology core for their assistance. We also thank Chun-Teng Huang and Chih-Cheng from the SBP Functional Genomics Core and John Naughton from the Salk GT3 Core for assistance with AAV production. Rebecca Porritt and Kang Liu from the Genomics and Spatial Transcriptomics Core provided assistance with library preparation and bulk RNA sequencing, and Svetlana Maurya from the Proteomics Core supported mass spectrometry sample processing and analysis. The research performed by the SBP Functional Genomics Core in this publication was supported by the Shared Instrumentation Grant S10 OD036254 and the NCI Cancer Center Support Grant P30 CA030199.

