## Supplementary Figures 1-4 for "Targeting CyclinD1-CDK6 to Mitigate Senescence-Driven Inflammation and Age-Associated Functional Decline"

**Supplementary Figure 1 | Validation of knockdowns and CDK4/6 inhibition in senescent IMR90s.**
a, qPCR validation of siRNA-mediated knockdown of CCND1, CDK4, and CDK6 in IR-induced senescent IMR90 fibroblasts. Expression normalized to the geometric mean of GAPDH and RPL13, shown as fold change relative to siNT controls.
b–c, Principal component analysis (PCA) of RNA-seq profiles for siRNA knockdowns (b) and Palbociclib vs DMSO-treated senescent cells (c), showing distinct clustering of CCND1/CDK6 knockdowns and Palbociclib-treated cells, respectively.
d, Co-immunoprecipitation of CCND1 followed by Western blot confirms association with CDK6 in senescent cells.
e–f, qPCR showing downregulation of SASP and ISG transcripts upon knockdown of CCND1 or CDK6, but not CDK4. Expression normalized to GAPDH and RPL13, and shown as fold change relative to siNT controls
g, Western blot analysis of Palbociclib-treated senescent IMR90s reveals reduced phospho-STAT1, total STAT1, and phospho-p65 levels, with unchanged total p65; Ponceau used as loading control.
Each biological replicate represents an independent irradiation. Error bars denote mean ± s.d. Statistical analysis performed using one-way ANOVA. *$P < 0.05$ considered significant.

**Supplementary Figure 2. Transcriptomic clusters and additional functional assays**
a, Heatmap of all differentially expressed genes (DEGs) across siRNA knockdown conditions. A red box highlights genes selectively upregulated with CCND1 or CDK6 knockdown, but not with CDK4 knockdown.
b, Hallmark pathway enrichment for the boxed cluster in (a).
c, Heatmap of cell cycle–related transcripts across siRNA knockdown conditions showing continued suppression, confirming maintenance of cell-cycle arrest.
d, Heatmap of all DEGs across Palbociclib versus DMSO-treated senescent IMR90s. A red box highlights genes upregulated with Palbociclib treatment.
e, Hallmark enrichment for the boxed cluster in (d).
f, Heatmap of cell cycle–related transcripts across Palbociclib versus DMSO conditions showing continued suppression, confirming maintenance of cell-cycle arrest.
g, Immunofluorescence quantification of CCF frequency and nuclear γH2AX intensity in senescent IMR90s after siRNA knockdown. Both phenotypes are reduced with CCND1 and CDK6 knockdowns compared to non-targeting controls.
h–i, Ranked bar plots from immunoprecipitation–mass spectrometry (IP–MS) using two independent CCND1 antibodies (h: antibody 1; i: antibody 2), showing top high-confidence CCND1 interactors in senescent IMR90s. Bars display sequence coverage and peptide counts for CCND1 IP versus IgG control.
Each biological replicate represents an independent irradiation. For immunofluorescence quantification, each dot reflects the average of ≥3 technical replicates per replicate.

**Supplementary Figure 3. Validation and long-term effects of Ccnd1 knockout in the aged liver.**
a, Immunohistochemistry staining of liver sections from young (4-month) and old (22-month) mice showing increased CCND1 protein in old livers, restricted to phospho–Histone H3–negative hepatocytes; partial hepatectomy samples included as proliferative controls.
b, Quantitative validation of *Ccnd1* editing efficiency in hepatocytes 3 weeks after AAV8-TBG-saCas9–mediated delivery of sgCcnd1 in 17-month-old mice. Indel formation and knockout efficiency were assessed using ICE (Inference of CRISPR Edits, Synthego).
c, qPCR analysis of ISG transcript levels following 3-month hepatocyte-specific *Ccnd1* knockout in aged (22-month) mice, showing reduced expression compared to age-matched Rosa26 KO controls. Expression values were normalized to the geometric mean of GAPDH and HPRT and are plotted as fold change relative to Rosa26 KO.
d, Western blot confirming sustained loss of CCND1 and reduced γH2AX, STAT1, and phospho-

STAT1 levels in aged livers after 3-month knockout.
e, Validation of γH2AX antibody specificity for detection of cytoplasmic chromatin fragments (CCFs). Irradiated young livers show increased nuclear γH2AX staining compared to unirradiated controls, confirming antibody performance in this context.
Each biological replicate represents an individual mouse. Error bars denote mean ± s.d. Statistical significance was assessed using the Mann–Whitney U test for qPCR and Welch's t-test for Western blots. *$P < 0.05$ was considered significant*

**Supplementary Figure 4. Ccnd1-positive hepatocytes exhibit a senescent-like transcriptional signature.**
a–b, Spatial transcriptomic (CosMx) analysis of young (4-month) and old (22-month) mouse livers showing increased expression of *Cdkn1a* and interferon-stimulated genes (ISGs) in *Ccnd1*-positive hepatocytes compared to *Ccnd1*-negative cells.
c–f, Single-cell RNA-sequencing analysis of liver datasets from the SenNet consortium, including young (3–4 months) and old (21–23 months) male and female mice. *Ccnd1*$^{+}$ hepatocytes show significant enrichment for both the SenMayo and ISG senescence gene signatures compared to *Ccnd1*$^{-}$ cells across both age groups, supporting a link between *Ccnd1* expression and a senescent transcriptional phenotype. Each biological replicate represents an individual mouse.

**Supplementary Figure 1**

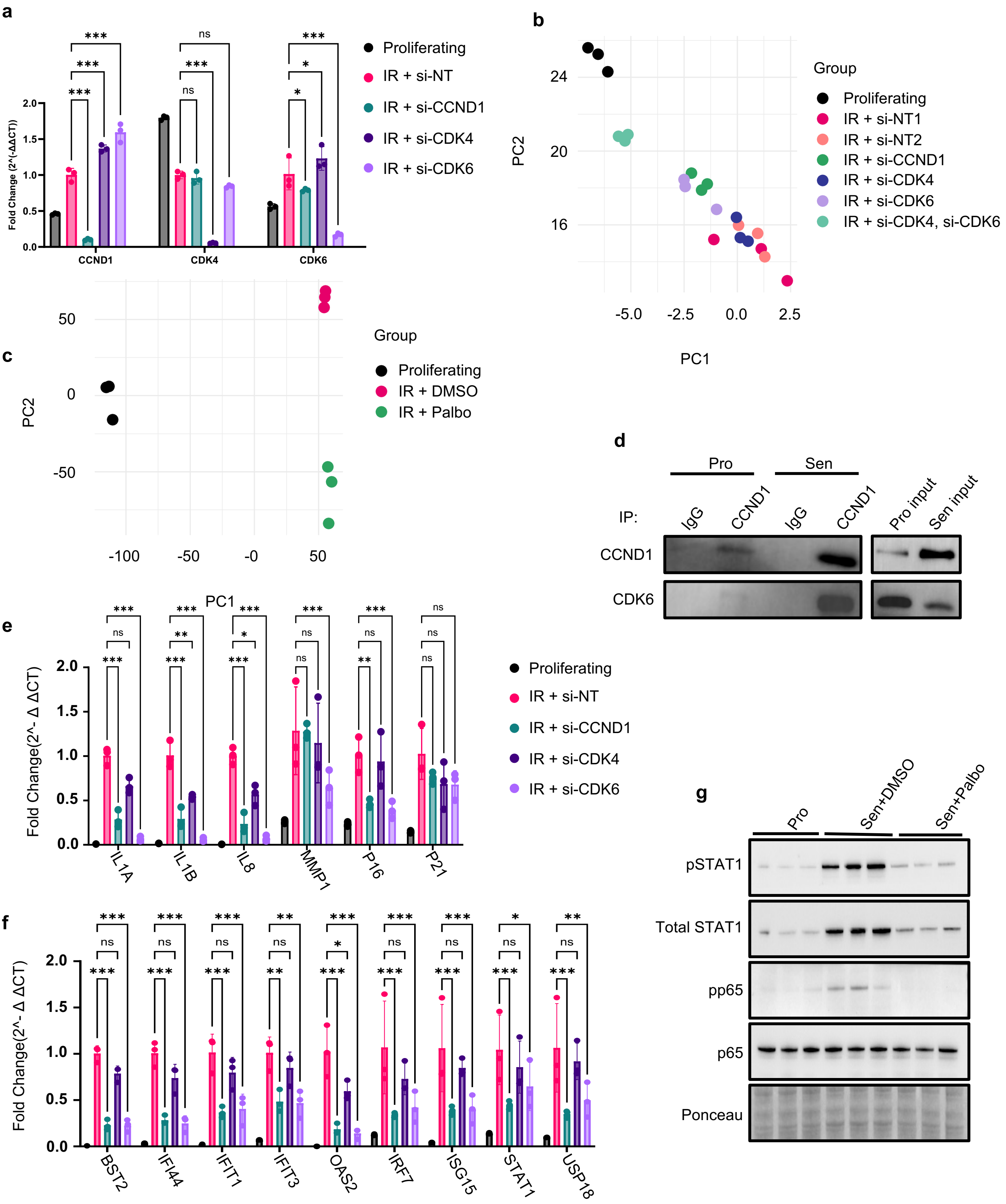

### Supplementary Figure 2

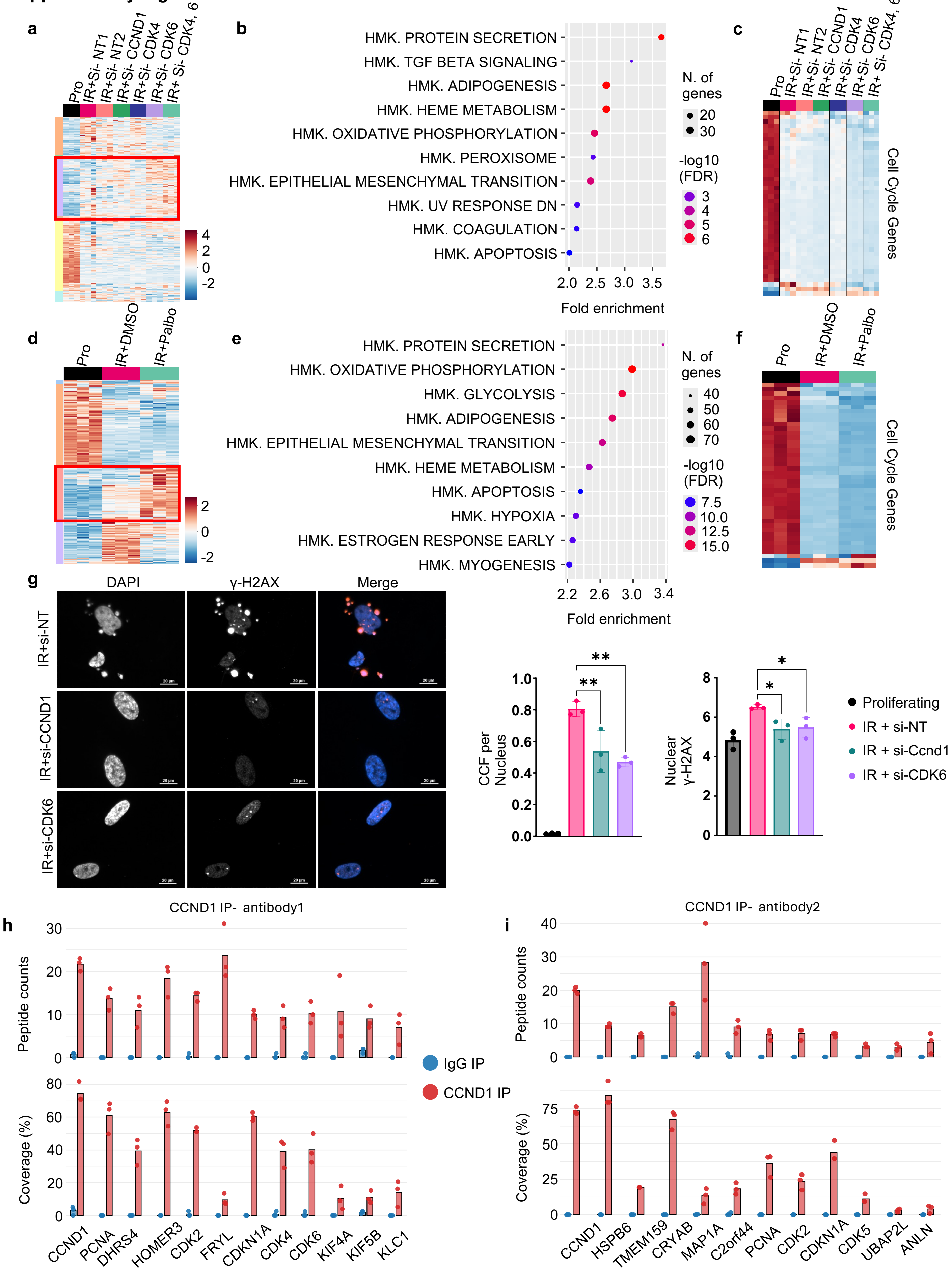

### Supplementary Figure 3

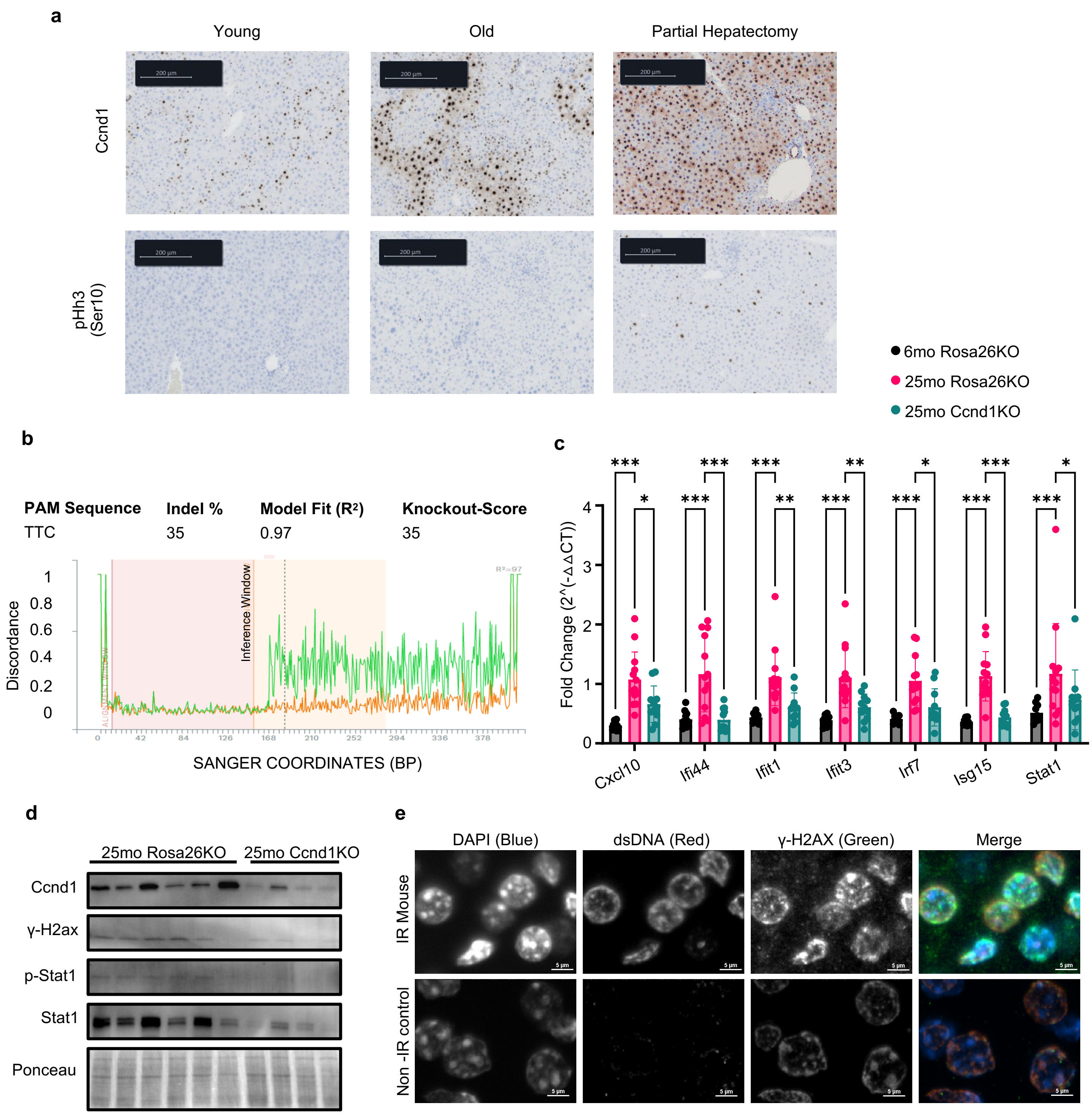

### Supplementary Figure 4

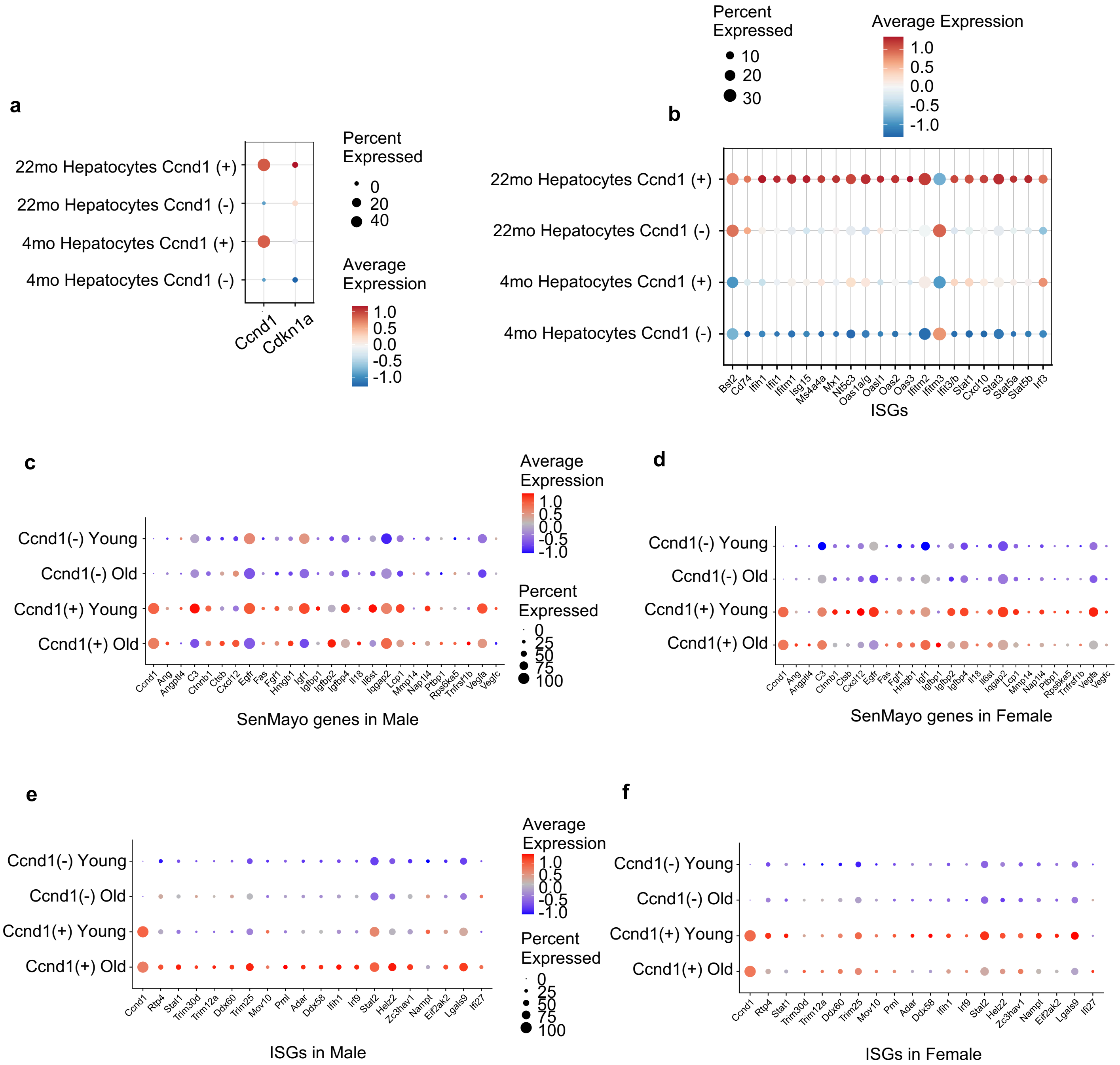
